# A microfluidics-assisted methodology for label-free isolation of CTCs with downstream methylation analysis of cfDNA in lung cancer

**DOI:** 10.1101/2022.02.28.482380

**Authors:** Ângela Carvalho, Catarina Guimarães-Teixeira, Vera Constâncio, Mariana Fernandes, Catarina Macedo-Silva, Rui Henrique, Fernando Jorge Monteiro, Carmen Jerónimo

## Abstract

Lung cancer (LC) is a major cause of mortality. Late diagnosis, associated with limitations in tissue biopsies for adequate tumor characterization contribute to limited survival of lung cancer patients. Liquid biopsies have been introduced to improve tumor characterization through the analysis of biomarkers, including circulating tumour cells (CTCs) and cell-free DNA (cfDNA). Considering their availability in blood, several enrichment strategies have been developed to augment circulating biomarkers for improving diagnostic, prognostic and treatment efficacy assessment; often, however, only one biomarker is tested. In this work we developed and implemented a microfluidic chip for label-free enrichment of CTCs with a methodology for subsequent cfDNA analysis from the same cryopreserved sample. CTCs were successfully isolated in 38 of 42 LC patients with the microfluidic chip. CTCs frequency was significantly higher in LC patients with advanced disease. A cut-off of 1 CTC/mL was established for diagnosis (sensitivity=76.19%, specificity=100%) and in patients with late-stage lung cancer, the presence of ≥ 5 CTCs/mL was significantly associated with shorter overall survival. MIR129-2me and ADCY4me panel of cfDNA methylation performed well for LC detection, whereas MIR129-2me combined with HOXA11me allowed for patient risk stratification. Analysis of combinations of biomarkers enabled the definition of panels for LC diagnosis and prognosis. Overall, this study demonstrates that multimodal analysis of tumour biomarkers via microfluidic devices may significantly improve LC characterization in cryopreserved samples, constituting a reliable source for continuous disease monitoring.

## INTRODUCTION

Lung cancer (LC) remains the leading cause of cancer-related mortality worldwide, with approximately 2.2. million new cases and 1.8 million deaths reported in 2020.^1^

Lung cancer is mostly categorized into two main histological groups: non-small cell lung cancer (NSCLC), originating from the lining epithelium, representing about 85% of all cases, and small cell lung cancer (SCLC), an aggressive is a less frequent neuroendocrine tumour, often presenting with metastatic disease and having few therapeutic options available.^2, 3^

Despite significant developments in LC management over the last decade, high mortality persists, mostly owing to the lack of early detection tools, entailing that most patients are diagnosed with late-stage disease. Furthermore, limitations in tissue biopsy availability and the absence widely implemented non-invasive tools to detect minimal residual disease (MRD) and early recurrence further impair more effective patient management. Lack of comprehensive knowledge on markers that may aid in early diagnosis, disease monitoring and identification of therapeutic targets also contribute to the high mortality rates. ^4–6^

Nevertheless, significant progress has been made recently, with the employment of minimally invasive techniques for cancer screening and monitoring, based on blood-derived cancer biomarkers, which may significantly impact on LC patient’ outcome and enable Precision Medicine. Indeed, the analysis of tumour biomarkers present in liquid biopsies (blood) and in bodily fluids (urine, pleural and cerebrospinal fluid) can provide real-time assessment of the disease and its molecular and genetic landscape. Biomarkers such as circulating tumour cells (CTCs), cell free DNA containing circulating tumor DNA (cfDNA, ctDNA), extracellular vesicles (EVs), tumour educated platelets (TEP) and cell-free RNAs have been widely studied over the last decade, with CTCs and ctDNA demonstrating clinical utility for prognostication and tumour mutational burden assessment. ^7–10^ Tumour biomarkers have also been widely investigated in several cancer progression pathways, uncovering their involvement and utility in processes such as metastasis formation, therapy resistance and potential therapeutic targets. ^11–13^ So far, CTCs and cfDNA remain the most widely investigated tumour biomarkers with most studies focusing on the independent analysis of a single biomarker. Recent reviews have addressed current research on single CTCs and cfDNA isolation and analysis in cancer.^14, 15^ However, CTCs and cfDNA complimentary nature may provide a more comprehensive analysis of LC, considering their single advantages and limitations. ^16^ CTCs are a key player in cancer progression, having a significant participation in the complex cancer metastatic process through the intravasation, circulation in the bloodstream and colonization of distant sites, forming metastasis. Thus, CTCs identification and quantification may be relevant for early detection, prognostication, and treatment monitoring. Isolated CTCs may be studied as whole cells, or through RNA, DNA and protein-based profiling.^17, 18^ CTCs have been shown to associate with LC progression and overall survival. ^10, 19^ ctDNA, on the other hand, represents a fraction of total cfDNA, which offers an invaluable opportunity for diagnosis and prognostication analysis through real-time detection of relevant genetic alterations such as point mutations and gene amplification, as well as epigenetic alterations like DNA methylation. Moreover, it may assist in therapy effectiveness monitoring, considering its quantitative and qualitive changes. cfDNA levels evaluation, as well as tracking of therapy efficacy by detecting acquired resistance mutations may allow for the assessment of disease progression, MRD, and early prediction of relapse.^20–23^ In LC, cfDNA has proven useful in the clinical setting for liquid biopsy-based analysis of EGFR mutations.^7^ Epigenetic alterations have also been increasingly evaluated in cfDNA and may assist in early diagnosis. Aberrant DNA methylation is commonly found in cancer and has been shown as a promising biomarker for diagnosis using cfDNA. DNA methylation at the promoter region of several cancer-related genes has been used to distinguish LC from benign conditions, and also to discriminate among LC subtypes.^23, 24^ The minute amount present in clinical samples and the accuracy required for cancer detection, led to the development of several strategies for efficient enrichment of biomarkers. Among them, microfluidics-based methodologies have stood out as particularly sensitive and efficient technologies for biomarker isolation and analysis. Application of microfluidic devices to liquid biopsies may enable low-cost, standardized, and automatized methodologies for cancer screening and monitoring in a high-throughput manner, applicable in the clinical setting. ^4, 25, 26^

Herein, we report the development and implementation of a microfluidic chip for CTCs isolation coupled w ith downstream analysis of cfDNA from the same sample. The system was validated in a retrospective cohort of LC patients, demonstrating potential for analysis of cryopreserved samples. Samples were processed through the microfluidic chip and label-free CTCs were isolated by physical properties. The remaining processed sample was used for cfDNA extraction and methylation analysis. CTCs were enumerated and phenotypically characterized via immunofluorescence and their diagnostic and prognostic value was assessed. Similarly, cfDNA methylation analysis was performed using a panel composed by HOXA11, MIR129-2 and ACDY4 gene promoters, aiming at LC detection and histological subtypes discrimination.

### Experimental section

#### Patient enrollment and sample collection

A retrospective cohort of LC patients was included in this study, comprising 42 patients diagnosed with LC between 2017 and 2021 at Portuguese Oncology Institute of Porto, Portugal, without previous oncological treatment. For control purposes, 32 blood samples donated between 2016 to 2021 by healthy volunteers were also analyzed.

Plasma was isolated by centrifuging whole blood in collected EDTA-containing tubes at 2500 rcf for 30 min at 4 °C and stored at – 80 °C until further use. Relevant clinical and pathological data was reviewed, and a clinical database was constructed.

This study was approved by the ethics committee of Portuguese Oncology Institute of Porto (CES-IPOPFG-EPE 177/018). All patients included in this study provided informed consent, in accordance with the Declaration of Helsinki ethical principles.

#### Microfluidic chip fabrication

The microfluidic system was designed in a CAD software (AutoCAD, Autodesk) and prepared using standard microfabrication techniques. Briefly, silicon (Si) photomasks with the microfluidic layout were prepared via photolithography in a clean room facility (Class 100). The microfluidic systems were then developed in polydimethylsiloxane (PDMS, Sylgard 184, Dow) by soft-lithography. PDMS was mixed with a curing agent at a ratio of 10:1 (wt/wt), degassed and poured into the Si mask. After baking at 60°C for 4h, the PDMS was peeled from the mask, cut and the inlets and outlet were punched. Finally, the PDMS layer with the microfluidic design was bonded to a glass slide with O_2_ plasma (Diener Zepto), developing the final, closed device.

#### Cell capture assay

Efficiency tests were conducted with MG63, a human bone osteosarcoma cell line (ATCC, USA). MG63 cells were cultured in 25 cm^2^ tissue culture flasks with high glucose Dulbecco’s Modified Eagle Medium (DMEM, Sigma Aldrich), supplemented with 10% fetal bovine serum (FBS, Gibco) and 1% Penicillin/Streptomycin (Sigma-Aldrich) in a humidified atmosphere of 5% CO_2_ at 37 °C.

Upon reaching confluence, cells were washed with PBS and detached by incubation with 0.5% (w/v) trypsin for 5 minutes at 37 °C, harvested and resuspended in culture medium. The concentration of cells in suspension was determined using a haemocytometer and 500 cells were spiked into 2 mL of cell culture medium. Prior to samples processing through the chip, a solution of 0.9 % sodium chloride (NaCl) was inserted into the chip at a flow rate of 200 μL/min to clear the system of air bubbles. The cell solution was pumped into the microfluidic chip at a flow-rate of 25 μL/min. Following isolation, cells were fixed with 4% paraformaldehyde (PFA) and fluorescently labelled for enumeration. Fixed cells were washed with NaCl and permeabilized with 0.1% Triton X-100 solution (v/v), for 10 min at room temperature (RT). Subsequently, a solution of blocking buffer (AbCAM) was loaded in the chip for 20 min. Cells were then stained with Alexa Fluor 488 (Life Technologies), at a 1:100 for 1 h, and washed with NaCl. Nuclei was stained with DAPI at 1:200 (Life technologies) for 10 minutes. Finally, cells were washed to remove excess dye and observed using an inverted fluorescence microscope (Axiovert 200M, Zeiss).

#### Clinical samples processing

Cryopreserved plasma samples (approx. 2 mL) were gently thawed at room temperature (RT) in the cryovial. Samples were mixed with 0.9 % NaCl (Sigma-Aldrich) at a 1:1 dilution and 5% of Ethylenediaminetetraacetic acid (EDTA, 0.4M, pH 8, Panreac), loaded into a 5 mL luer-lock syringe and pumped into the microfluidic chip using a syringe pump (DARWIN microfluidics). The system was placed in an inverted microscope (Leica) and connected to the syringe and the collection tube with Tygon tubings and luer-lock connectors (Darwin Microfluidics). The samples were introduced into the chip at a flow rate of 25 μl/min, with a total processing time of 80 minutes for a 2 mL sample. The remaining sample was transferred to a collection tube and used for further analysis of the cfDNA present in plasma.

#### In-chip immunofluorescence detection of isolated CTCs

Isolated CTCs were fluorescently labelled inside the microfluidic chip. Cells were fixed with 4% paraformaldehyde for 10 min at RT, permeabilized with 0.1% triton X-100 (Sigma-Aldrich) and incubated with blocking buffer solution (AbCAM) for 20 min. CTCs were labelled with mouse anti-human CD326 for NSCLC (EpCAM, Biolegend) and mouse anti-human CD133 for SCLC at 1:100 (Biolegend), followed by goat anti-mouse IgG H&L Alexa Fluor 647 at 1:400 (AbCAM), Alexa fluor 594 anti-Vimentin at 1:200 (Biolegend) and rabbit anti-human Alexa fluor 488 anti-CD45 (AbCAM) at 1:400. Cells were washed with NaCl, and nuclei stained with DAPI (Life Technologies) at 1:200 for 10 minutes. Finally, stained cells were washed with NaCl for excess dye removal and observed using an inverted fluorescence microscope (Axiovert 200M, Zeiss).

#### Circulating-free DNA Extraction, sodium-bisulfite modification and preamplification

Following samples processing via the microfluidic chip, circulating cell-free DNA was extracted from 400 μL of plasma using the MagLEAD 12gC extractor (Precision System Science Co.) with the MagDEA DX SV kit according to the equipment’s protocol and eluted in 50 μL. cfDNA was stored at −20°C until further use. Then, 20 μL of each extracted cfDNA sample and 5μg/20μL of Human HCT116 DKO Non-Methylated DNA and Human HCT116 DKO Methylated DNA were sodium-bisulfite-modified using EZ DNA Methylation GoldTM Kit (Zymo Research) according to the manufacturer’s recommendation. Bisulfite-modified DNA was eluted in 10 μL of sterile distilled H2O and stored at −80 °C until further use. SsoAdvancedTM PreAmp amplification was performed in 8 μL of the sodium-bisulfite-modified DNA following manufacturer’s recommendations. The preamplified DNA was diluted in 50 μL of sterile distilled H_2_O, in a final volume of 100 μL, and stored at −20°C until further use.

#### Multiplex qMSP

Promoter methylation levels of three genes (MIR129-2me, ADCY4me and HOXA11me) were evaluated by singlepex and multiplex quantitative methylation-specific PCR (qMSP), using the preamplified DNA as template. The housekeeping gene *ACTβ* was used as an internal reference gene to normalize the assay. For each gene, primers and TaqMan probes with specific fluorochromes and quenchers (Table 1) were wielded. All lung cancer samples were run in triplicate in 384-well plates using an Applied BiosystemTM QuantStudioTM 12K Flex Real-Time PCR System. Standard curves in each plate were used with six serial dilutions (5x factor dilution), allowing for relative quantification and PCR efficiency evaluation. Two wells of sterile distilled water were used as negative control in all plates. The multiplex gene panel including MIR129-2me, ADCY4me and ACTβ worked on 60 °C annealing temperature, while HOXA11me on 64 °C. All plates displayed efficiency values above 90% and relative methylation levels were defined as the ratio between the mean methylation levels of each gene and the respective value for *β-Actin*, multiplied by 1000 for easier tabulation.

**Table 1.**
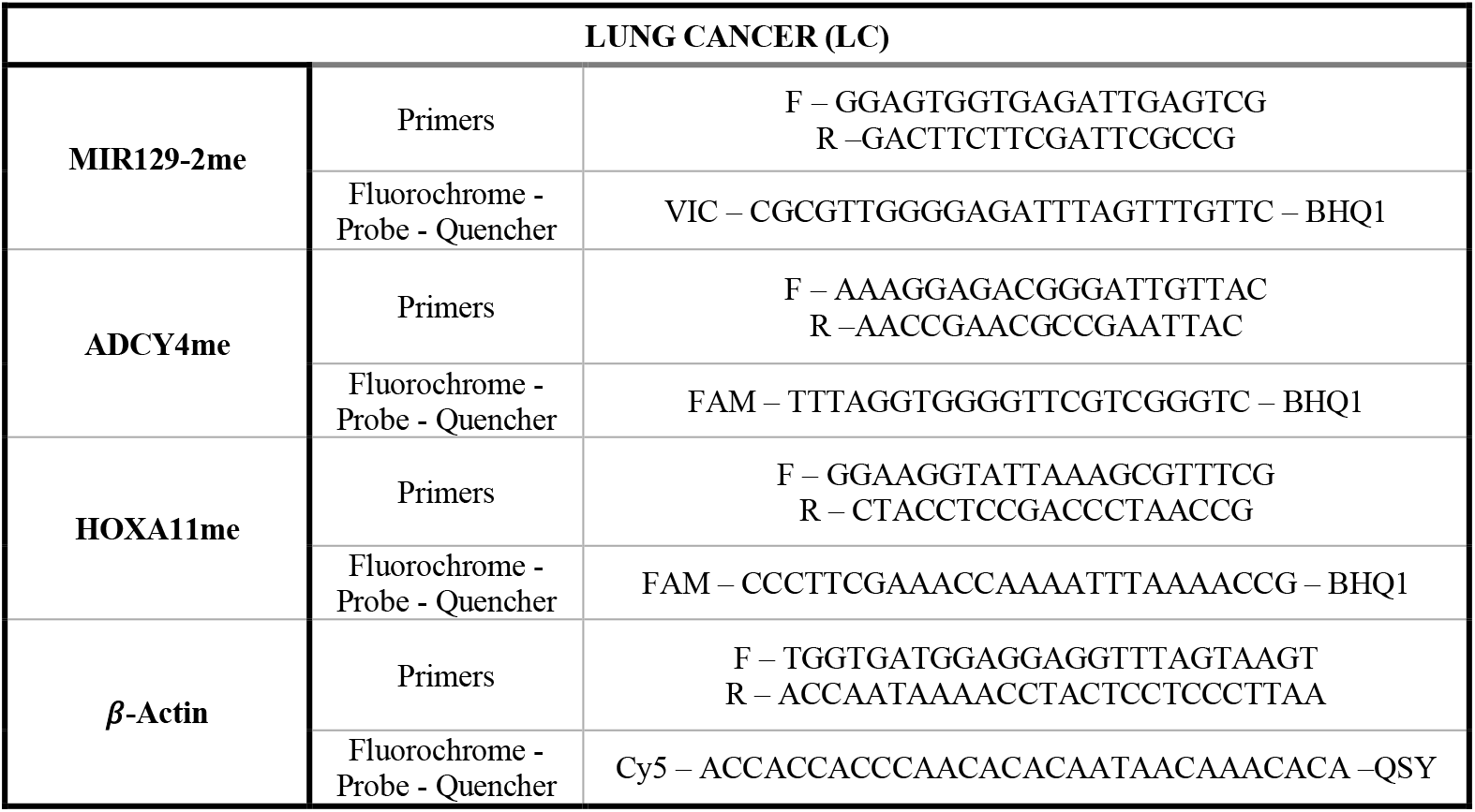
Primers and probes sequences with respective fluorochrome and quencher.

#### Statistical Analysis

Mann-Whitney U test was used for comparisons between two groups, while Kruskal-Wallis test was used for multiple groups, followed by Mann-Whitney U test with Bonferroni’s correction for pairwise comparisons. Spearman non-parametric test was performed to assess correlations between methylation levels and patient age, as well as ACTβ and cfDNA quantity levels. Receiver Operator Characteristic (ROC) curve analysis was performed to calculate the areas under the curve (AUC). Validity estimates (sensitivity, specificity) with 95% confidence intervals were determined for CTCs biomarker performance. Kaplan-Meier curves were constructed, and log-rank test was used to compare overall survival (OS) between groups, considering clinicopathological variables and CTCs enumeration. OS was calculated as the time between the date of diagnosis and the date of patient death. A backwards multivariable Cox-regression model comprising all significant variables on univariable analysis was computed to determine whether CTCs levels were independently associated with OS. A result was considered statistically significant when p-value ≤0.05. The statistical analysis was performed using the GraphPad Prism 9.0.0 and IBM SPSS statistical software’s.

## RESULTS AND DISCUSSION

### Patients clinicopathological characterization

A total of 42 LC patients and 32 healthy volunteers were included in this study. The selected 32 healthy volunteers were matched for LC patients age range. Clinical data from patients is shown in table 2.

**Table 2.** Clinicopathological features of the LC patients enrolled in this study.

| Patients (n= 42) |  |  |  |  |
| --- | --- | --- | --- | --- |
| Age median (range) |  |  | 68 (44-83) |  |
| Gender | Male |  | 69 (64-83) |  |
|  | Female |  | 64 (44-80) |  |
| Clinicopathological features |  |  |  |  |
| Tumor stage | Early | I | 21 (50%) | 16 (38%) |
|  |  | II |  | 5 (12%) |
|  | Late | III | 21 (50%) | 9 (21%) |
|  |  | IV |  | 12 (29%) |
| Smoking habits | Smoker |  | 33 (79%) |  |
|  | Non-smoker |  | 9 (21%) |  |
| Metastasis | Yes |  | 17 (40%) |  |
|  | No |  | 25 (60%) |  |

### Microfluidic system cell capture efficiency

The designed microfluidic chip is composed by three isolation chambers with sequential rows of micropatterns of decreasing interspacing for label-free isolation of CTCs (Figure 1 A). Each chamber contains 850 micropatterns with interspacing varying from 50 μm to 15 μm. Each chamber discloses two filtration areas. The initial rows of micropatterns gap varies between 50 and 20 μm and were designed for bulk filtration, formed by squared patterns in triangular disposition which can held cell-clusters or small clogs that may occur and through which single cells can flow. The final rows of the isolation area consist of micropatterns with sizes between 40μm x 40μm and 100μm x 40μm, and interspacings from 20μm to 15μm, aiming at single cell isolation. CTCs isolation was performed under laminar flow, with a flow rate of 25 μl/min, ensuring a cellular trajectory that could maintain cell integrity during the processing. The flow rate was defined following initial tests with fluorescent PMMA particles (PolyAn GmbH) and MG63 cells and assessment using an optical microscope. MG63 cells were chosen for assessing system efficiency based on the fact that mean size lays within the range of larger CTCs (approximately 18 μm diameter).^27^ Cell spiking assays with MG63 cell line revealed a cell isolation efficacy of 74%.

**Figure 1.**
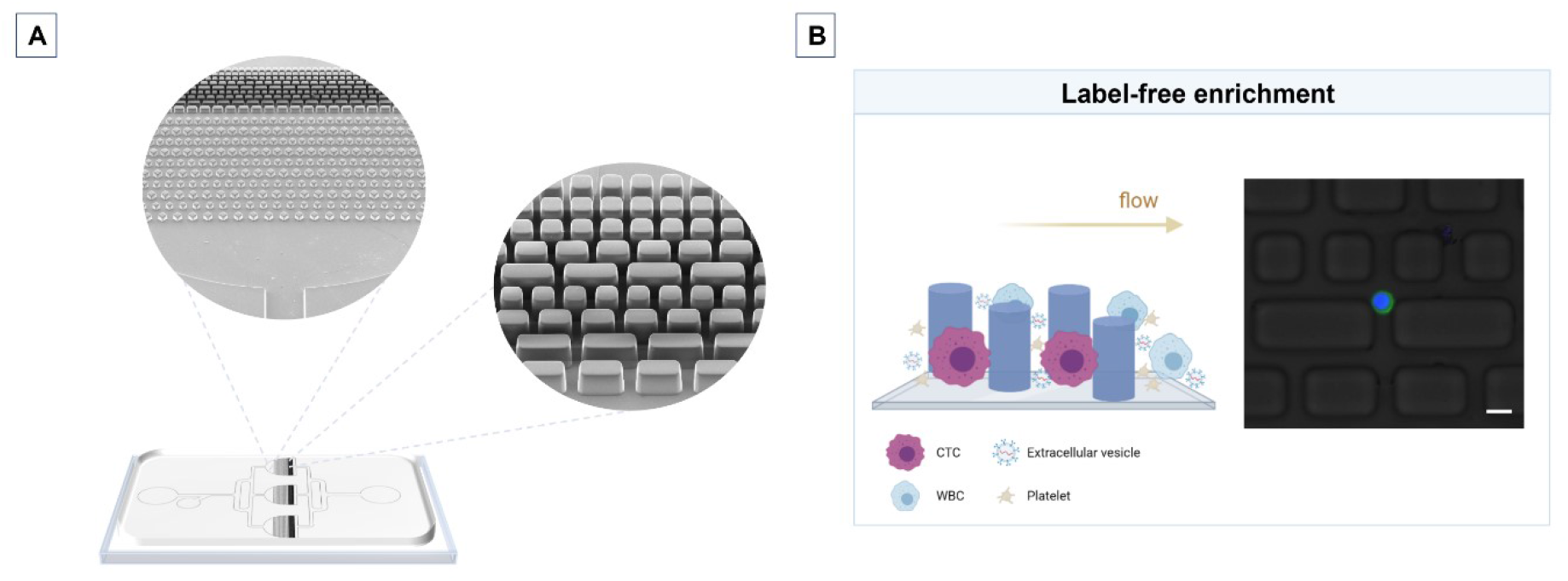
Microfluidic system design and patterns arrangement in the isolation chambers (A) and label-free principle for cells enrichment and isolated MG63 cell in the single cell isolation area of the microfluidic device (B). Scale bar: 20 μm

The microfluidic device was designed for capturing larger CTCs and to diminish leukocyte contamination to increase CTCs purity for molecular analysis, although it may also reduce the system efficiency in capturing smaller cells. Indeed, the observed size range for captured cells was higher than the patterns gap (> 15 μm). (Figure 1B).

### CTCs detection via immunofluorescence

Plasma samples from 42 LC patients and 32 healthy donors were processed using the developed microfluidic chip. Following isolation in the microfluidic chip, CTCs were observed and counted by immunofluorescence staining. CTCs were distinguished from white blood cells (WBC) using CD45 to stain WBC and epithelial, mesenchymal and EMT markers to stain CTCs (EpCAM/CD133/VIM). Cells positive for at least one of the CTCs markers (EpCAM+/CD133+/VIM+) and DAPI+, and negative for CD45 (CD45-) were considered CTCs. Features such as round shape, high nuclear to cytoplasm ratio and a minimum size of 4 μm also allowed for CTCs identification. ^28^ CD45+/DAPI+ cells were identified as WBCs (Figure 2).

**Figure 2.**
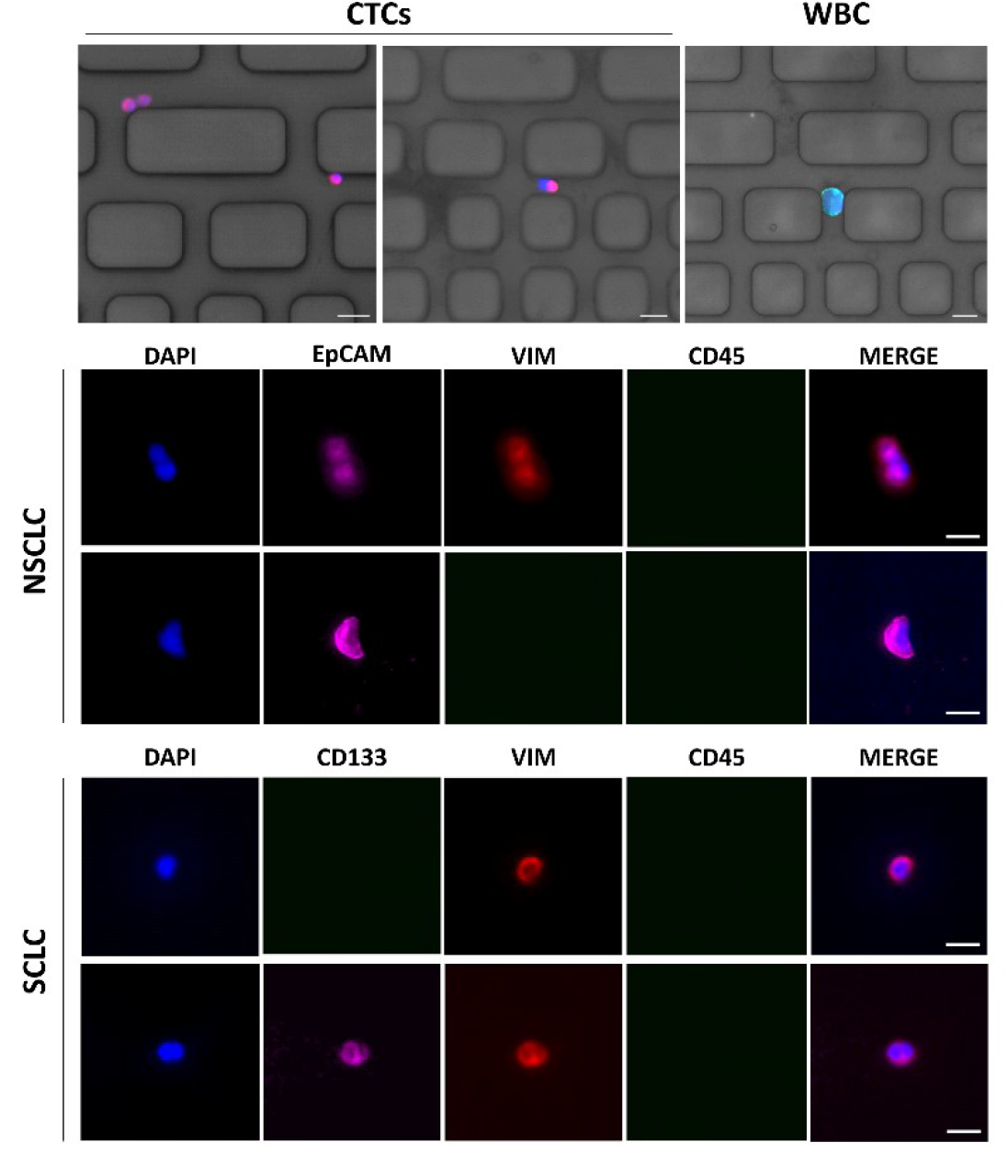
Representative images of phenotypical heterogeneous circulating tumour cells (CTCs) detected in non-small cell lung cancer (NSCLC) and small cell lung cancer (SCLC) patients, and a white blood cell (WBC) isolated in the microfluidic device. Scale bars: 20 μm

CTCs were captured between micropatterns in the isolation area based on their size. The single cell isolation area of the microfluidic system maximized cell-substrate interaction due to the ranging sizes of the micropatterns and contributed to more efficient CTC capture, when associated with the defined flow rate. Additionally, adhesion to the microstructures also allowed for CTCs isolation (Figure 2). Indeed, plasma proteins adsorption to the micropatterns might have increased CTCs adhesion. Cancer cells display increased adhesion capacities when compared to normal blood cells. Chen *et al* making use of this property, developed a nanoroughness microfluidic CTC capture chip that displayed a superior capture capacity of heterogeneous CTCs populations.^29, 30^

The CTCs antibody-independent isolation method allowed for the capture and characterization of CTCs with heterogeneous phenotypes. Following enrichment, NSCLC CTCs were stained with EpCAM and VIM, while SCLC were stained with CD133 and VIM. EpCAM was expressed in 56.3% of all identified NSCLC CTCs, whereas VIM was expressed in 76.2 % NSCLC CTCs. Both surface markers were expressed in a total of 43.75 % samples. EpCAM, an epithelial cell adhesion molecule, is typically expressed by epithelial cancers.^31^ In fact, EpCAM is a surface marker used in the only method approved by the Food and Drug Administration (FDA) for CTCs isolation in colon, breast and prostate cancer, CellSearch (VERIDEX). As for LC patients, although tested in different studies, it has not received approval, yet.^32, 33^ While EpCAM analysis guarantees a fair comparison with CellSearch, EpCAM-based detection also presents limitations. Specifically, cancer cells that undergo epithelial to mesenchymal transition (EMT), facilitating invasion and dissemination by entering in the circulation, display low EpCAM levels, whereas mesenchymal markers might be expressed. Indeed, CTCs adopting EMT typically display characteristics of cancer stem cells (CSCs), including higher migratory potential to assist in invasion, self-renewal and differentiation.^34^ In the past few years, Vimentin (VIM) was suggested as a robust marker of CTCs undergoing EMT, being expressed in CTCs from several different cancers.^35–37^ In NSCLC, CTCs’ Vim expression has been also recognized and associated with metastasis development and worse prognosis. ^38, 39^

Recently, Xie *et al* demonstrated that cell surface vimentin (CSV) might be considered a LC biomarker, disclosing 0.67 sensitivity and 0.87 specificity, using a cut-off of 2 CTCs per 7.5 mL of blood. CSV^+^ CTCs were identified in 83.33% of LC patients and correlated with cancer stage, lymph node involvement and distant metastasis. Additionally, the team reported that CSV^+^ CTCs showed a better diagnostic performance than serum tumour makers, such as carcinoembryonic antigen (CEA), neuron-specific enolase (NSE), cancer antigen 125 (CA125), and CA153.^40^ Thus, the high number of VIM^+^ CTCs isolated in NSCLC patients further advocates the need to apply EMT markers for CTCs accurate detection.

Other authors have also observed higher CTC detection by using methods that isolate heterogeneous CTCs populations. Krebs *et al* previously reported a higher CTCs count in NSCLC using ISET, a label-free method and the successful enrichment of CTCs negative for epithelial markers, including EpCAM. ^41^ EpCAM was also shown to be downregulated in CTCs and tissue of lung neuroendocrine tumours.^42^

For further marker validation, SCLC CTCs were identified as CD133+ and/or VIM^+^. CD133 was expressed in 70% of all analysed SCLC samples, while VIM^+^ cells were found in 80% of the cases. Both markers were found in 50% of all identified CTCs. SCLC is a neuroendocrine tumour commonly associated with rapidly progressive disease and dismal prognosis. CD133 has been previously tested as LC surface marker for cells with higher tumorigenic potential and stem-like characteristics and used as a potential biomarker for SCLC CTCs. ^43, 44^ Interestingly, Sarvi *et al* observed a correlation between CD133 expression and SCLC chemoresistance, as well as high tumorigenicity *in vitro* and *in vivo*. Moreover CD133^+^ cells were found to have higher neuropeptide receptor expression, displaying sensitivity to a novel neuropeptide antagonist, peptide-1, highlighting a potential utility of selective targeting of chemoresistant cells, thus providing a novel therapeutic opportunity for resistant SCLC. ^45^

VIM expression in SCLC has also been associated with increased dissemination potential and metastatic activity, entailing reduced OS. ^46^ Among the 10 SCLC patient samples analysed, 7 were CD133^+^ and 8 were VIM^+^, showing that both surface markers can provide utility in SCLC CTCs enumeration and confirming the higher tumorigenic potential and aggressiveness of SCLC. However, further analysis with a larger patient cohort is necessary for proper validation.

Multimarker immunofluorescence analysis of CTCs has been increasingly applied for CTCs detection, covering CTCs heterogeneity, and improving CTCs identification, while specific LC CTCs markers are not validated. Figure 2 shows representative images of CTCs isolated in the microfluidic chip and their phenotypic heterogeneity.

### Diagnostic and prognostic value of CTCs in LC

In 42 LC patient samples processed in the microfluidic chip, CTCs were detected in 90.5% (n=38) of all samples, comprising 81% of early-stage (17/21) and 100% of late-stage (21/21) LC patients. Furthermore, 50% of patients disclosed 4 or more CTCs/mL. Remarkably, quantification revealed significant differences among stage I, II and stage III, IV patients, with increasing CTCs count in late-stage samples, as expected (Figure 3A).

**Figure 3.**
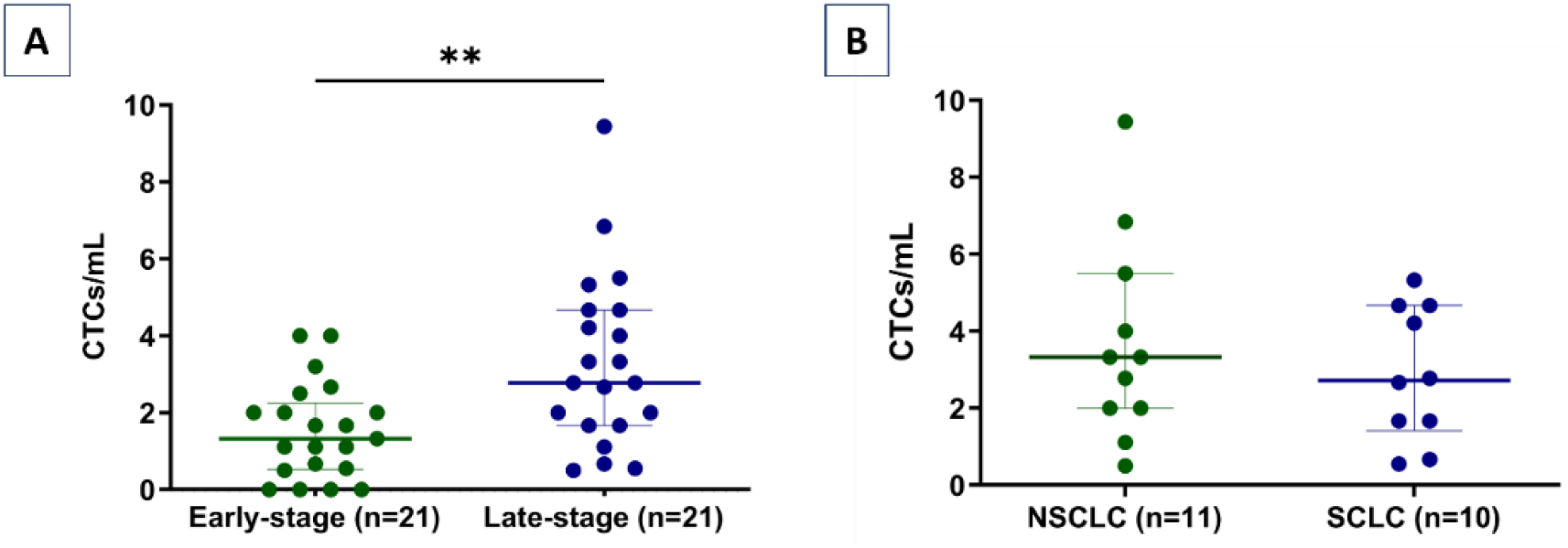
CTCs enumeration plots for (A) early-stage and late-stage LC patients and (B) Non-small cell lung cancer (NSCLC) and small cell lung cancer (SCLC). Lines represent the interquartile range and median value. ** represents statistically significant differences between groups (Mann-Whitney, p < 0.01)

Indeed, early-stage patients displayed a median of 1 CTC/mL (range: 0-4 CTCs/mL), while samples of late-stage cases presented a median of 3 CTCs/mL (range: 1-9 CTCs/mL). CTCs were not detected in any of the 32 healthy individuals of the control group. Complete leukocyte depletion was achieved in 84% of the processed samples. Aggregates of 2 cells were identified in 3 patients but not considered CTCs clusters as these are typically composed of 3 or more CTCs. Other authors have reported on successful microfluidic-based approaches for CTCs isolation in LC. Zhou *et al* described a multi-flow microfluidics platform (MFM) for CTCs isolation in NSCLC. The device showed a recovery rate efficiency of between 87% and 93% in assays performed with cancer spiked cells and CTCs detection in 6 out of 8 stage IV NSCLC samples, with a median of 12 CTCs per mL of whole blood.^**47**^ On the other hand, the Parsortix™ Cell Separation System (ANGLE), a semi-automated microfluidic based technology for CTCs isolation through size and deformability, has been applied for label-free enrichment of CTCs in SCLC patients. Cytokeratin (CK) positive CTCs were detected in all 12 patients enrolled in the study, with an average number of recovered CTCs >20 CTCs.^48^

CTCs enrichment and detection techniques influence CTCs enumeration as well as sample processing methods. Depending on the blood’s volume, sample preparation protocols, isolation method and CTCs classification criteria, there is significant variability of results. Therefore, a careful analysis considering the methodologies and parameters applied in each study is essential. Herein, we focused on the analysis of 2 mL cryopreserved plasma samples. In addition to a low sample volume processed *in-chip*, the cryopreservation of the liquid biopsy samples might have contributed to lower cell frequency, as freezing and thawing may affect cell viability. Nonetheless, Brungs *et al* demonstrated that cryopreserved blood samples were a good source for gastroesophageal CTCs isolation and characterization. However, significantly lower CTCs number was found comparing cryopreserved and fresh blood samples, which particularly relevant in samples with high CTC count (>50), and less significant for samples with low CTC count. Moreover, CTCs isolated from cryopreserved samples were found to be an independent prognostic factor for OS. Importantly, the duration of cryopreservation did not affect CTC numbers. ^**49**^ While cryopreserved samples may disclose lower frequency of CTCs, they may still provide invaluable clinical utility when paired with efficient methods for CTCs isolation and more sensitive analysis techniques (ddPCR, single-cell analysis), allowing for tumour analysis via liquid biopsies whenever clinicians deem necessary. Other influencing factors for CTCs frequency are the pre-analytic sample preparation protocols. The samples used in our study were previously centrifuged, with removal of RBCs layer, where some part of the PBMC layer may be lost, also contributing to a lower CTC number and leukocyte contamination. In addition, the designed interspacing of the microfluidic system might also affect CTCs enumeration. Albeit small CTCs identified in the system adhered to the microstructures, the design aimed at larger CTCs isolation and effectively allowed for a significant leukocyte depletion while smaller CTCs may have been lost. Indeed, a size-based isolation approach may enhance CTCs enumeration and improve phenotypic characterization when compared with EpCAM-based isolation strategies.

Indeed, higher CTC quantification was reported by others using label-free methodologies.^50–52^ Importantly, in our hands and in accordance with other reports on CTCs enumeration in LC, CTCs quantification displayed prognostic value for disease aggressiveness and allowed for patient stratification, with statistically significant higher CTC rates in late-stage disease. ^53^ The diagnostic performance of CTCs in LC was also assessed. When CTCs cut-off was set to 1 CTC/mL, we were able to identify LC with 76.19% sensitivity and 100% specificity, an AUC of 0.952. CTCs performance for stage discrimination revealed a 61.90% sensitivity and 80.95% specificity for late-stage detection with a cut-off of 2.5 CTCs/mL (AUC: 0.763). In patients with late-stage disease, no significant differences for CTCs count were observed between NSCLC (n=11) and SCLC (n=10). Remarkably, 8 SCLC and 10 NSCLC patients displayed metastatic disease (Figure 3B). In general, SCLC patients exhibit significantly higher CTCs values compared to other LC patients, with increased CTCs numbers correlating with worse overall survival (OS) and progression free survival (PFS). ^54, 55^

However, in our hands, NSCLC displayed slightly higher median CTCs (3.3 CTCs/mL) than SCLC (2.7 CTCs/mL). Both patients with the highest CTCs counts in NSCLC (total count: 17 and 13 CTCs) displayed significantly shorter OS than all the remaining patients, with an OS lower than 1 month. Hence, although further validation is required, increased CTCs frequency associates with disease aggressiveness.

### Correlation of CTCs Frequency with LC Patients’ Clinicopathological data

High CTC count was associated with poor OS in a diverse range of cancers. In our study, the prognostic value of CTC/mL for OS of LC patients was analysed with Kaplan-Meier survival curves for the whole patient cohort (n=42) and late-stage patients (n=21) (Figure 4). CTCs cut-off was defined as 3 CTCs/mL for the patient cohort and was associated with significantly reduced OS for patients ≥ 3CTCs/mL. The mean OS for patients with less than 3 CTCs/mL at diagnosis was 841 days (95% CI, 690-992) compared with 448 days (95% CI, 191-705) for patients with CTC counts ≥ 3 CTCs/mL. In late-stage patients, a specified CTC cut-off of 5 CTCs/mL associated with significantly reduced OS. Patients with < 5 CTCs/mL displayed a mean overall survival of 431 days (95% CI, 254-609), while patients with ≥ 5CTCs/mL showed 136 days mean OS (95% CI, 0-282). Univariate analysis in late-stage patients showed that cell counts ≥ 5CTCs/mL was significantly associated with shorter OS. In multivariate Cox regression analysis of all LC patients, CTCs number did not retain prognostic value after adjustment for stage, cancer subtype and presence of metastasis. Several studies have investigated the prognostic value of CTCs to determine OS in LC patients with a wide range of CTCs cut-offs. ^53^ Indeed, among 125 stage IIIB-IV NSCLC patients ≥5 total CTCs per 7.5 mL of peripheral blood associated with significantly reduced OS but not progression-free survival (PFS).^56^ In NSCLC, Hofman and colleagues defined 50 CTCs/10 mL at baseline to be independently associated with shorter OS. ^57^ Also, similarly to our results, Nieva et al defined 5 CTCs/mL as a predictor of shorter OS. ^58^ Cheng et al defined 10 CTCs/7.5 mL as an optimal cut-off for OS prediction in 91 SCLC patients with extensive disease at baseline. Patients with <10 CTCs at baseline disclosed significantly improved median OS. ^59^ Moreover, Hou et al demonstrated that ≥50 CTCs/7.5 mL independently associated with reduced OS in SCLC patients ^60^, while Igawa et al reported ≥2 CTCs/7.5 mL to be independently associated with OS in all SCLC patients stages. ^61^ As previously discussed, differences in samples’ processing protocols, CTCs enrichment methods, sample type and CTCs cut-off contribute to disparate results obtained in the several studies published so far. However, in line with previous reports, CTC quantification strongly associated with survival, emphasizing its value for LC prognostication.

**Figure 4.**
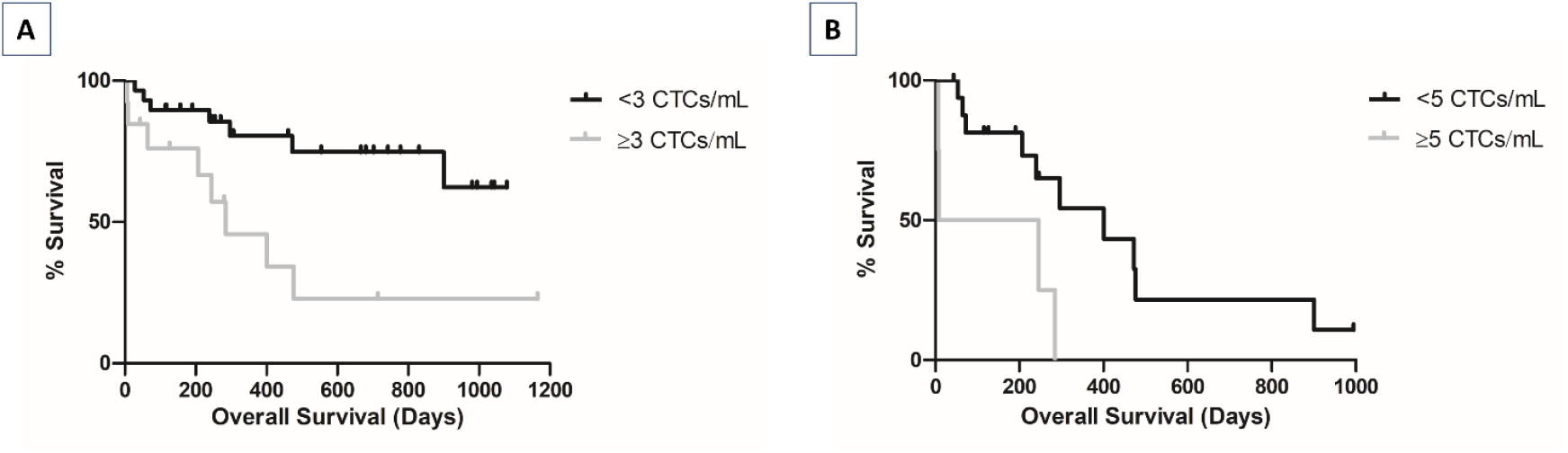
Kaplan-Meier overall survival predictive value of CTC count in all LC patients (A) and late-stage LC patients (B). Kaplan–Meier curves evaluated in the whole cohort of patients with CTCs cut-off value ≥ 3 CTCs/mL (Log-rank test, p = 0.007) and in advanced stage III–IV patients, with a CTCs cut-off value ≥ 5 CTCs/mL (Log-rank test, p=0.017).

### cfDNA methylation

#### Lung Cancer Prediction, Stage and Histological Subtypes

cfDNA is a well-known promising biomarker, under study for diagnostic and precision therapy purposes^62^. Because aberrant DNA methylation of cancer-related genes develops very early during tumorigenesis, its evaluation might be used for LC detection and subtyping ^23, 63–65^. Microfluidics have been increasingly applied in liquid biopsies and may improve cancer biomarkers extraction and enrichment from biological fluids. They might also allow for combinatorial analysis of circulating biomarkers due to their unique processing parameters, not requiring reagents for specific isolation of a single biomarker. Therefore, cfDNA methylation using multiplex qMSP was evaluated following a protocol involving microfluidic-assisted filtration of circulating tumor cells. A total of four genes-ACTβ (reference gene) and MIR129-2, HOXA11 and ADCY4 (target genes)-were selected and tested for their potential in LC detection and screening.

cfDNA’s MIR129-2me (p= 0.0471) and ADCY4me (p= 0.0375) levels were significantly higher in LC patients’ cell-free DNA’s compared to controls. LC patients with advanced disease displayed significantly higher MIR129-2me (p= 0.0025) and HOXA11me (p= 0.0005) levels compared to patients with stage I / II disease. In line with previous studies, HOXA11me associated with advanced-stage LC ^66–68^. MIR129-2me (p= 0.0380), HOXA11me (p= 0.0023) and ADCY4me (p= 0.0465) levels were significantly higher in SCLC patients comparing with NSCLC patients (Figure 5). Furthermore, when analyzing only NSCLC patients, the highest MIR129-2me (p= 0.0299) and HOXA11me (p= 0.0460) levels were also found in patients with advanced stage disease (Figure 6).

**Figure 5.**
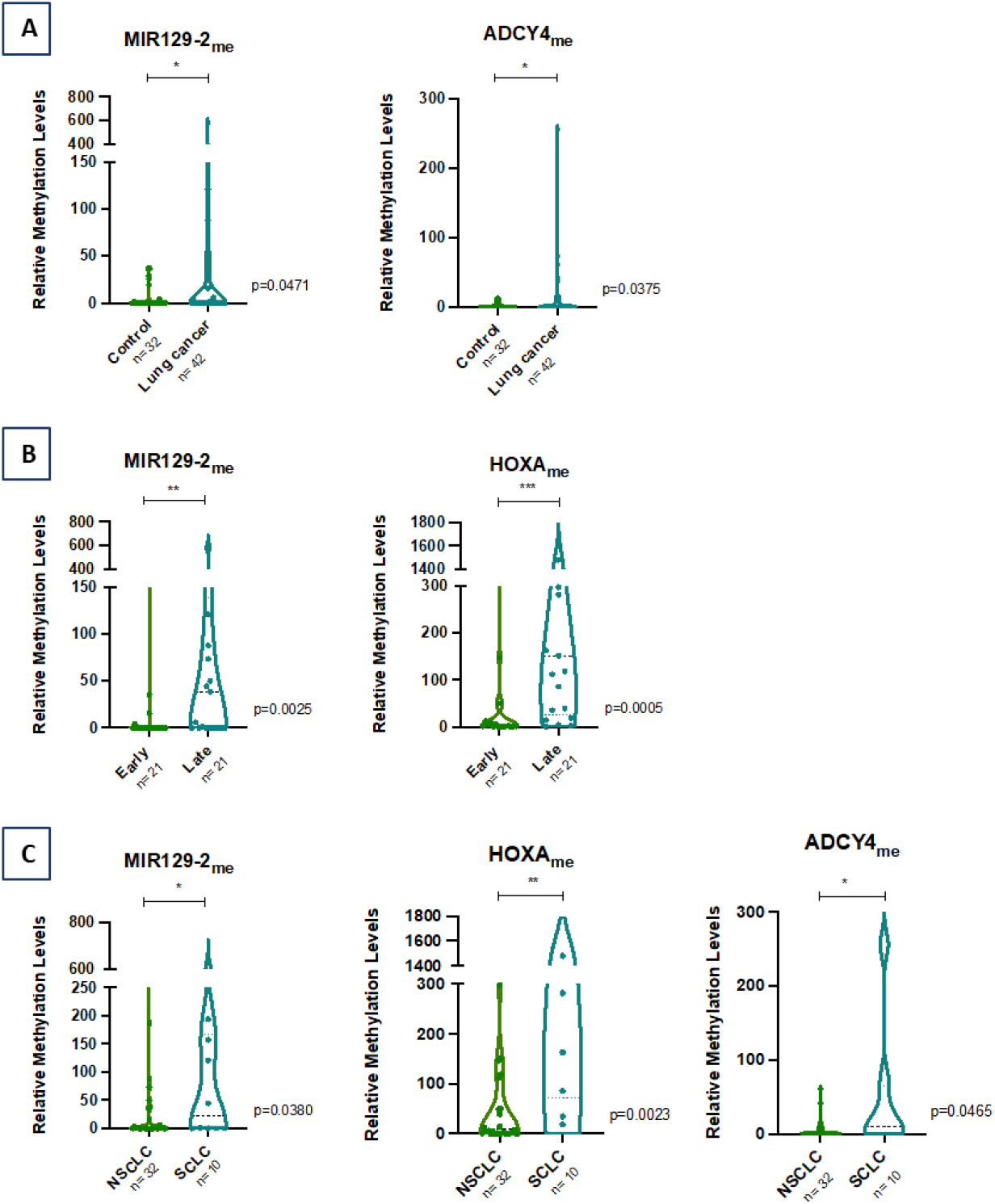
Distribution of MIR129-2me, HOXA11me and ADCY4me relative methylation levels in (A) Controls (n = 32) and LC samples (n = 42); (B) Early-stage (n= 21) and late-stage (n= 21); (C) Histological group NSCLC (n=32) and SCLC (n= 10). Mann-Whitney U Test, n.s. p > 0.05, *p < 0.05, **p < 0.01, ***p < 0.001, ****p < 0.0001.

**Figure 6.**
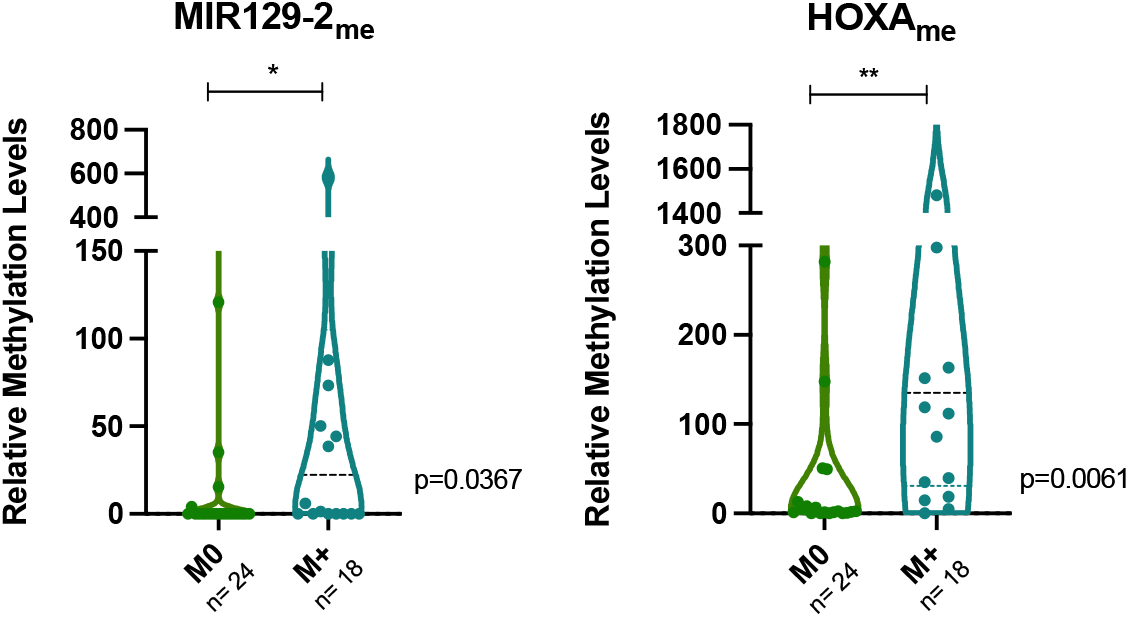
Distribution of MIR129-2me and HOXA11me relative methylation levels with NSCLC in Early-stage (n= 21) and late-stage (n= 11). Mann Whitney U Test, n.s. p > 0.05, *p < 0.05, **p < 0.01, ***p < 0.001, ****p < 0.0001.

Likewise, significantly higher circulating MIR129-2me (p= 0.0367) and HOXA11me (p= 0.0061) levels were found in LC patients with distant metastases (Figure 7). Nonetheless, no significant associations were found for other clinicopathological variables.

**Figure 7.**
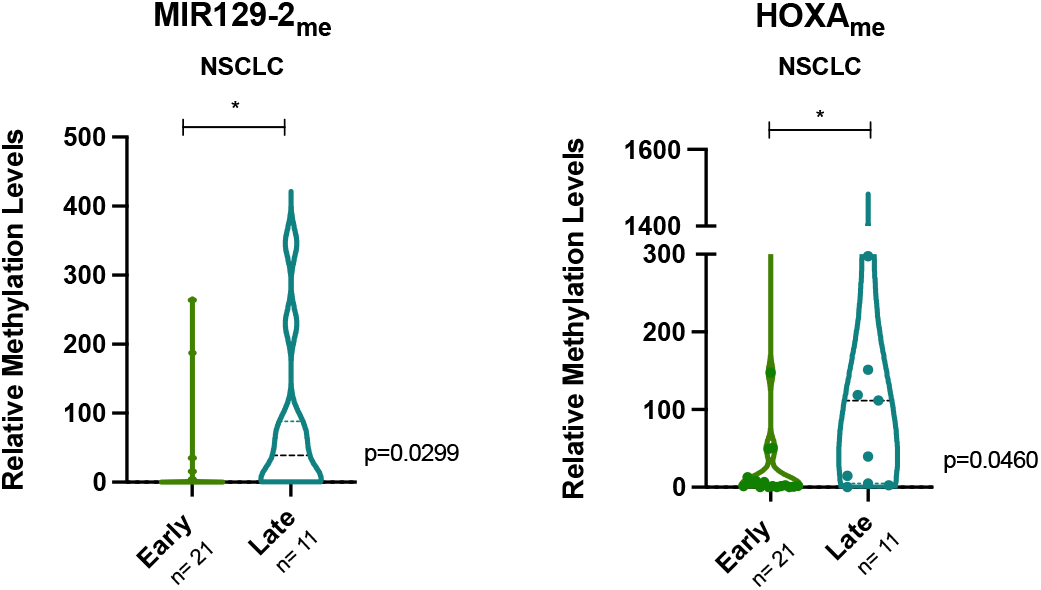
Distribution of MIR129-2me and HOXA11me relative methylation levels according with metastatic status. Mann Whitney U Test, n.s. p > 0.05, *p < 0.05, **p < 0.01, ***p < 0.001, ****p < 0.0001.

Interestingly, circulating MIR129-2me (p= 0.0282) levels were significantly higher in smokers vs non-smokers patients (Supplementary Figure 1). This is in line with previous reports on AHRR and F2RL3 genes ^69^.

### Biomarker performance of cfDNA methylation

Concerning biomarker performance of methylation-based cfDNA processed by a microfluidic device, ADCY4me showed the highest sensitivity in detecting LC, while HOXA11me displayed the highest specificity (100% and 93.75%, respectively) (Table 3). Interestingly, ADCY4me was previously shown to identify breast cancer in plasma samples ^70^.

**Table 3.** Biomarker performance of each gene promoter methylation for LC detection in cfDNA.

| Gene | AUC<br>(95 %<br>CI) | Sensitivity<br>% (95%<br>CI) | Specificity<br>% (95%<br>CI) | Accuracy<br>% (95%<br>CI) |
| --- | --- | --- | --- | --- |
| MIR129-2me | 0.572 | 29.55 | 87.50 | 53.95 |
| HOXA11me | 0.563 | 29.55 | 93.75 | 56.58 |
| ADCY4me | 0.577 | 100 | 34.38 | 72.37 |

Remarkably, combined in a panel (MIR129-2me, ADCY4me and HOXA11me), the highest accuracy and sensitivity for LC detection was obtained when at least 1 gene was positive, whereas the highest specificity (96.88%) was reached when 3 genes were positive (Table 4).

**Table 4.** Biomarker performance detection of panel (MIR129-2me, HOXA11me and ADCY4me) in cfDNA.

|  | At least 1<br>positive | At least 2<br>positive | All 3<br>positive |
| --- | --- | --- | --- |
| **Sensitivity % (95% CI)** | 100 | 47.62 | 11.90 |
| **Specificity % (95% CI)** | 31.25 | 87.50 | 96.88 |
| **Accuracy % (95% CI)** | 70.27 | 64.86 | 48.65 |

Furthermore, the performance of the two biomarkers to discriminate early-stage from late-stage disease was also evaluated. MIR129-2me and HOXA11me displayed the highest specificity and sensitivity, respectively, for identification of late-stage LC (90.48%% and 85.71%, respectively). The panel composed of MIR129-2me and HOXA11me with at least 1 positive biomarker disclosed a sensitivity of 90.48%, while positivity for both biomarkers depicted 100% specificity for late-stage disease identification (Table 5).

**Table 5.** Biomarker performance distinguish early-stage and late-stage patients of panel (MIR129-2me and HOXA11me) in cfDNA.

|  | At least 1 positive | All 2 positive |
| --- | --- | --- |
| **Sensitivity % (95% CI)** | 90.48 | 47.62 |
| **Specificity % (95% CI)** | 57.14 | 100 |
| **Accuracy % (95% CI)** | 73.81 | 73.81 |

### Multimodal biomarkers performance

Then, we assessed the diagnostic and prognostic value of combined CTCs and cfDNA methylation in LC, in cryopreserved plasma samples. The highest performance for LC detection was achieved combining CTC and at least one methylated promoter, depicting 76.19% sensitivity and 100% specificity, matching the performance of single CTCs analysis (1 CTC/mL) and further improving the specificity for LC detection achieved with cfDNA methylation panel with 1 positive gene (31.25%).

Importantly, the assessment of both biomarkers was able to identify late-stage patients with a positive CTCs detection (cut-off: 2.5 CTCs/mL) or 1 positive gene methylation with a sensitivity of 100% and a specificity of 75%. When comparing with the single biomarker’s performance, increased sensitivity was obtained comparing with single CTCs analysis (61.90%) and 1 positive gene in the methylation analysis panel (90.48%). Multimodal biomarkers analysis showed improved specificity comparing cfDNA methylation (57.14%), although lower than single CTCs (80.95). Overall, combined analysis of both biomarkers revealed improved diagnostic performance for all lung cancer cases, particularly those with advanced stages.

Moreover, the combined assessment of both biomarkers provided additional disease characterization via cell count and surface markers identification and molecular characterization of LC. In this cohort, 4 early-stage patients did not have detectable CTCs, for whom cfDNA methylation analysis displayed at least 2 positive genes and enabled detection through the assessment of ADCY4me, MIR129-2me and HOXA11me panel, with accurate subtype discrimination. Likewise, CTCs were found in all three patients with only one promoter testing positive gene.

The establishment of a methodology for multimodal assessment of both biomarkers may close the gap on limitations associated with the analysis of a single biomarker.

While combinatorial assessments of multiple cancer biomarkers in a single sample are still scarce, some authors have reported on the high value of these analysis. Liu et al developed a workflow to detect EGFR mutations in cfDNA and CTCs in blood of 24 NSCLC patients. The combinatorial assessment resulted in non-invasive EGFR mutation analysis and provided preliminary validation of the applied workflow of a “Total Liquid Biopsy”. ^71^

A multimodal liquid biopsy based on CTCs and ctDNA assessment for the early monitoring and outcome prediction of chemoresistance in metastatic breast cancer was reported. CTCs and ctDNA were assessed at baseline and after four weeks of first-line chemotherapy. The team demonstrated a multivariate prognostic model including CTC count at four weeks (≥5CTC/7.5 mL), ctDNA variant allele frequency (VAF) at baseline, tumour subtype and grade. ^72^

Similarly, our analysis focused on combinatorial potential of CTCs and cfDNA for LC diagnosis, stratification and prognostication. CTC count and cfDNA methylation analysis provided evidence regarding the utility of different tumour biomarkers for a more complete LC diagnosis and patient personalized monitoring.

## CONCLUSIONS

Liquid biopsies can play an extraordinary part in the improvement of lung cancer patients screening and management, especially in cases where tissue biopsy cannot be performed. New technologies, such as microfluidic devices may be critical for the development of highly sensitive and efficient methodologies for cancer detection and characterization.

A critical advantage of the implemented methodology described in this work is the ability to perform sequential analysis of both biomarkers, allowing for a combinatorial assessment of different tumour characteristics, while eliminating sample-based bias. CTCs can provide insight regarding disease development with prognostic value and allowing for protein and nucleic acids screening, while cfDNA can be applied for analysis of epigenetic markers or mutational profiles for early diagnosis, monitoring and analysis of the disease mutational profile.

The microfluidic system accomplished CTCs enrichment in 90.5% of samples, with a low sample dilution and volume and the designed workflow subsequently allowed methylation analysis of cfDNA, establishing markers for LC detection and prognosis. Combinatorial assessment of both biomarkers was successfully attained in cryopreserved samples and established panels for LC detection (sensitivity: 76.19% and specificity: 100%) and prognostication (sensitivity: 100% and specificity: 75%). These can be of extreme importance when typical tumour analysis cannot be performed in the clinical setting or complementary analysis are needed. This methodology can improve patient personalized monitoring and be performed at any time during their follow-up. Some optimizations to the protocol should follow, with the validation of the system with fresh samples of increased volumes, smaller gap for size-based isolation, several genes analysis and an increased patient cohort, which may provide increased specificity and sensitivity performance of both biomarkers.

Overall, this study demonstrated the value of a chip-based methodology for complementary CTCs and cfDNA analysis as diagnostic and prognostic LC biomarkers, by means of minimally invasive liquid biopsies. More comprehensive liquid biopsies based on combined analysis of CTCs and cfDNA may be of high clinical relevance for LC diagnosis and monitoring.

## ACKNOWLEDGEMENTS

This work was supported by FEDER funds through COMPETE (POCI-01-0145-FEDER-030831) and by Portuguese funds through Fundação para a Ciência e a Tecnologia (PTDC/BTM-TEC/30831/2017) in the framework of project TRIMARKCHIP. V.C. received the support of a fellowship from “la Caixa” Foundation (ID 100010434). The fellowship code is LCF/BQ/DR20/11790013.

