## Supplemental Figure 1 for "A microfluidics-assisted methodology for label-free isolation of CTCs with downstream methylation analysis of cfDNA in lung cancer"

### Electronic Supplementary Information

#### for

a. i3S-Instituto de Investigação e Inovação em Saúde, Universidade do Porto, Rua Alfredo Allen 208, 4200-135 Porto, Portugal
b. INEB-Instituto de Engenharia Biomédica, Universidade do Porto, Rua Alfredo Allen 208, 4200-135 Porto, Portugal
c. Porto Comprehensive Cancer Centre (P.CCC), R. Dr. António Bernardino de Almeida, 4200-072 Porto, Portugal
d. Cancer Biology and Epigenetics Group, IPO Porto Research Center (GEBC CI-IPOP), Portuguese Oncology Institute of Porto (IPO Porto), R. Dr. António Bernardino de Almeida, 4200-072 Porto, Portugal
e. Department of Pathology, Portuguese Oncology Institute of Porto (IPO Porto), R. Dr. António Bernardino de Almeida, 4200-072 Porto, Portugal
f. Department of Pathology and Molecular Immunology, School of Medicine and Biomedical Sciences, University of Porto (ICBAS-UP), Rua Jorge Viterbo Ferreira 228, 4050-513 Porto, Portugal
g. Faculdade de Engenharia, Departamento de Engenharia Metalúrgica e Materiais, Universidade do Porto, Rua Dr Roberto Frias, s/n, 4200-465 Porto, Portugal

#### Corresponding author

* Ângela Carvalho


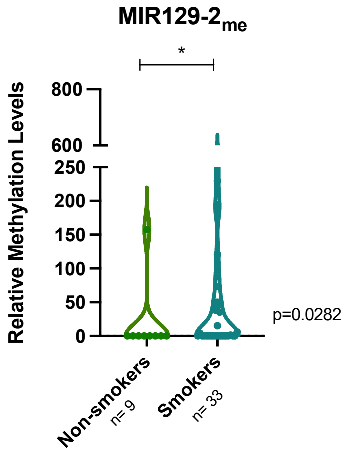


**Figure S1 - Distribution of MIR129-2me relative methylation levels according with smoking habits. Mann Whitney U Test, n.s. p > 0.05, *p < 0.05, **p < 0.01, ***p < 0.001, ****p < 0.0001.**
